# Bioaffinity-based surface-immobilization of antibodies to capture endothelial colony-forming cells

**DOI:** 10.1101/2021.06.23.449631

**Authors:** Mariève D. Boulanger, Mohamed A. Elkhodiry, Omar S. Bashth, Gaétan Laroche, Corinne A. Hoesli

**Author notes:** Co-first authors (Mariève D. Boulanger, Mohamed A. Elkhodiry).

## Abstract

Maximizing the re-endothelialization of vascular implants such as prostheses or stents has the potential to significantly improve their long-term performance. Endothelial progenitor cell capture stents with surface-immobilized antibodies show significantly improved endothelialization in the clinic. However, most current antibody-based stent surface modification strategies rely on antibody adsorption or direct conjugation via amino or carboxyl groups which leads to poor control over antibody surface concentration and/or molecular orientation, and ultimately bioavailability for cell capture. Here, we assess the utility of a bioaffinity-based surface modification strategy consisting of a surface-conjugated cysteine-tagged protein G molecules that immobilize Immunoglobulin G (IgG) antibodies via the Fc domain to capture circulating endothelial colony-forming cells (ECFCs). The cysteine-tagged protein G was grafted onto aminated substrates at different concentrations as detected by an enzyme-linked immunosorbent assay and fluorescence imaging. Different IgG antibodies were successfully immobilized on the protein G-modified surfaces and higher antibody surface concentrations were achieved compared to passive adsorption methods. Surfaces with immobilized antibodies targeting endothelial surface proteins, such as CD144, significantly enhanced the capture of circulating ECFCs *in vitro* compared to surfaces with non-endothelial specific antibodies such as anti-CD14. This work presents a potential avenue for enhancing the clinical performance of vascular implants by using covalent grafting of protein G to immobilize IgG antibodies more effectively.

**Table of Contents:** 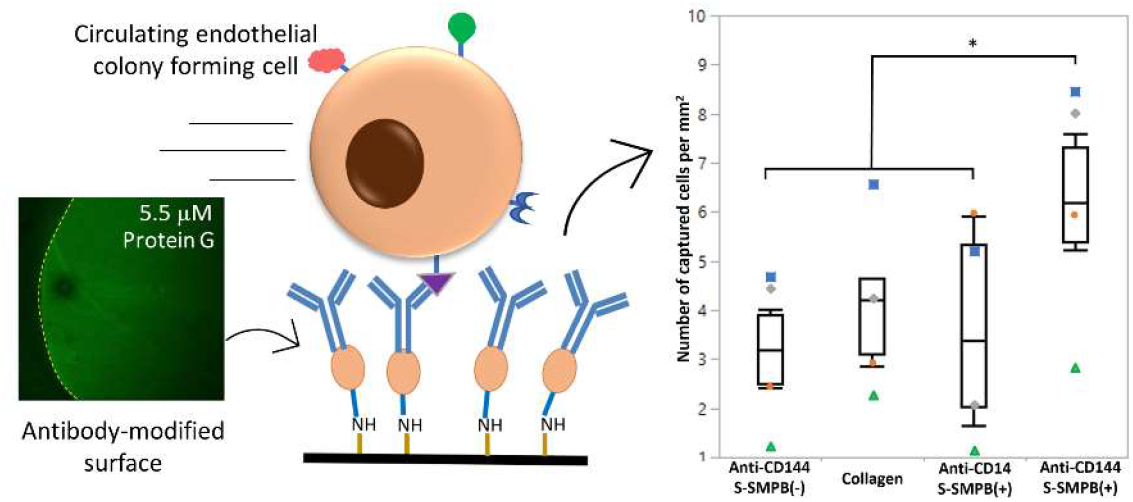

Antibody immobilization via surface-conjugated recombinant cysteine-protein G provides an effective approach to capture circulating therapeutic cells.

## Introduction

Despite decades of continuous improvement in the overall performance of vascular implants, biocompatibility challenges remain a major area of concern^1, 2^. A healthy endothelium can modulate protein deposition, platelet activation, and proliferation of the underlying smooth muscle layer which are critical in reducing the risk of re-stenosis and thrombosis^3^. A promising approach to enhance stent endothelialization and reduce the risks of implant failure is to capture circulating endothelial progenitor cells (EPCs)^4^. Capturing EPCs, particularly their functional subtype endothelial colony forming cells (ECFCs), accelerates the formation of a neo-endothelium due to their high proliferative potential and clonal expansion^5, 6^.

One way to promote ECFC capture on the surface of blood contacting devices is to modify the surface with antibodies that target ECFC surface antigens^7^. The surface-immobilized antibodies mimic the function of glycoproteins present on the vessel lining such as selectins and intercellular adhesion molecule-1 in the endogenous recruitment of circulating ECFCs to tissues undergoing vascular regeneration^8–10^. This antibody-based strategy was commercially adopted to create the Genous™ stent (Orbusneich, USA), an anti-CD34 antibody surface-modified metal stent. In a pilot study with 193 patients, the Genous™ stent was shown to be as safe as drug-eluting stents (the control group) with no observed difference in adverse cardiac effects. The study also revealed a promising reduction in the rate of in-stent thrombosis in patients with the Genous™ stent compared to the drug-eluting stent group despite an increased rate of re-stenosis^11^.

Since the first generation of EPC capture stents, potential areas of improvement in stent design were investigated. For example, the choice of CD34 as a target antigen has been under scrutiny due to its presence on the surface of hematopoietic progenitors that can exacerbate intimal hyperplasia^12^. Stents modified with anti-vascular endothelial cadherin (CD144) were shown to be more effective in accelerating endothelialization in animal models^13^. Furthermore, available antibody-modified implants mostly utilize passive adsorption or covalent conjugation to immobilize antibodies on the surface. Using passive adsorption, non-covalent interactions between the antibody and the surface dictates the strength and longevity of the surface modification often leading to lack of durability and lack of control over antibody orientation^14^. Using covalent conjugation, most strategies exploit reactive groups introduced on surfaces to conjugate antibodies via free amine or carboxyl groups present in the antibody sequence^15, 16^. The abundance of these functional groups in an antibody results in random antibody orientations on the surface and could lead to partial denaturation affecting its antigen binding efficacy^17, 18^.

More recently, improved bio-affinity-based antibody immobilization techniques emerged as an alternative that can enhance the potential of a new generation of ECFC-capturing stents^19–21^. Li *et al.* demonstrated that stent surfaces modified with anti-CD34 antibodies immobilized via the Fc-binding protein A improved *in vivo* stent endothelialization compared to unmodified controls^22^. However, due to the layer-by-layer coating strategy applied in this study, protein A molecules can take different conformations on the surface which reduces the availability of sites available for antibody immobilization and hence antibody surface density. Recombinant DNA technology can be used to introduce cysteine residues in the sequence of Fc-binding proteins which can then be conjugated onto surfaces via the thiol group. Gold substrates modified with cysteine-tagged protein G increased antibody surface density compared to adsorption controls while maintaining optimal antibody orientation leading to enhanced antigen binding capacity. This technique has been successfully applied to the design of immune-assays on a chip^23, 24^. Oriented surface immobilization of antibodies via covalent grafting of cysteine-tagged protein G remains untested for *in vivo* cell capture applications, particularly EPCs.

Here, we grafted cysteine-tagged protein G on polystyrene, the most commonly-used tissue culture plastic, in order to immobilize EPC capture antibodies. We hypothesized that our modified surfaces would increase ECFC capture compared to conventional antibody immobilization via passive adsorption. To test this hypothesis, a parallel plate flow chamber was used to study how ECFCs – derived from human donors – are captured under dynamic flow conditions.

## 1. Materials and Methods

### 2.1 Surface Modification and Antibody Immobilization

The bioaffinity-based antibody immobilization strategy relied on a 3-step process consisting of the activation of an aminated surface with an amine to sulfhydryl heterobifunctional linker which is subsequently used to graft Cys-protein G – through sulfhydryl/maleimide reaction – followed by adding IgG antibodies that are immobilized via their Fc region on the protein G molecules (Figure 1). First, aminated surfaces (Corning™ PureCoat™ Amine Culture Dishes, Thermo Fisher Scientific™) were reacted for 2 h with 150 μL/cm^2^ of a 3 mg/mL suspension of sulfo-succinimidyl-4-(*p*-maleimidophenyl)-butyrate (S-SMPB, #BC24, G-Biosciences) in phosphate buffered saline solution (PBS, #21600010, Thermo Fisher Scientific). Next, protein G was attached to the linking arm by adding 150 μL/cm^2^ of a 5.5 μM recombinant Cys-protein G (protein G with an N-terminal Cys residue added to the recombinant protein sequence, #PRO-1328, Prospec-Tany Technogene Ltd) suspension in PBS for 1 h. Finally, primary antibodies targeting cell surface antigens (mouse anti-human CD31 antibody #303101; mouse anti-human CD105 #323202; mouse anti-human CD144 #348502; and mouse anti-human CD14, anti-CD14, #367102; all from Biolegend, San Diego, US) were immobilized on the protein-G modified surfaces by adding 150 μL/cm^2^ of antibody solution at 5 μg/mL in PBS for 1 h. Surfaces were then rinsed twice with PBS, once with a 1% SDS-TRIS pH 11 solution (5% v/v of 20% sodium dodecyl sulfate #05030, from Sigma Aldrich and 2.4 % w/v TRIS base PBP151-500 from Fisher Scientific in reverse osmosis water, pH adjusted to 11 with 2N NaOH solution) to remove adsorbed molecules, twice with PBS, and finally rinsed with reverse osmosis (RO) filtered water. The surfaces were then air-dried and stored for at most 1 week at room temperature before use. Adsorption controls followed the same surface modification scheme, except that surfaces were not activated with S-SMPB prior to incubation with Cys-Protein G.

**Figure 1:**
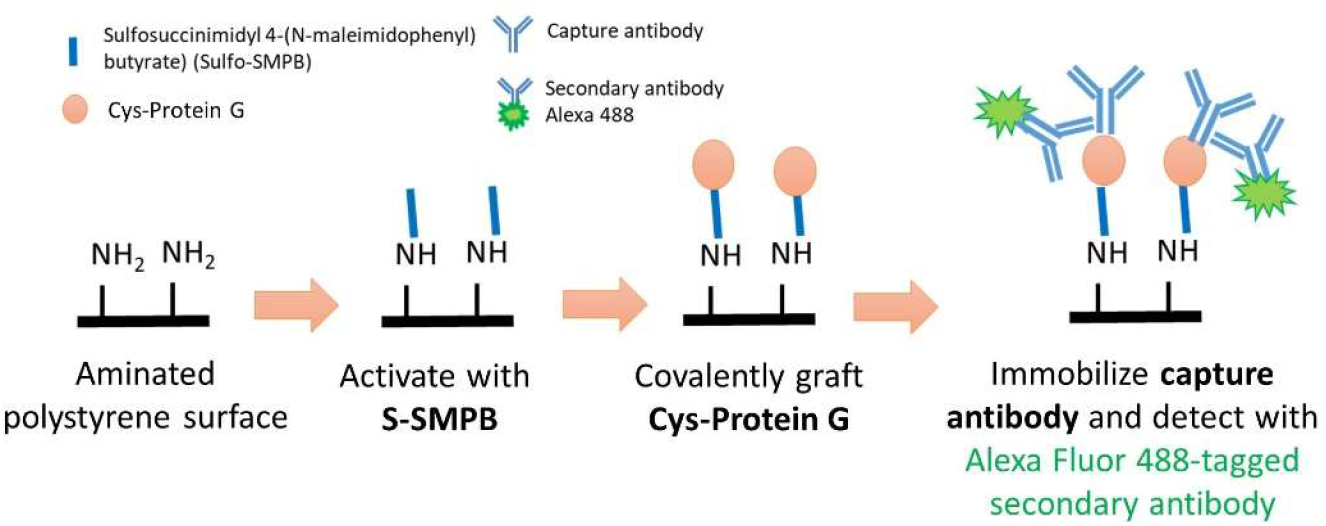
Schematic representation of the antibody immobilization process followed by a fluorescence-based antibody detection step.

For cell capture experiments under flow, surfaces were obtained by cutting aminated polystyrene Petri dishes (BD Purecoat™ Amine #354732, BD Biosciences, San Jose, USA) into 3.0 cm × 2.5 cm slides using a Micro Mill (Datron Neo 3-axis CNC Mill, Cell imaging and analysis network, McGill University, Canada). The circumference of these cut samples was lined with Teflon™ tape (#3213-103, polytetrafluoroethylene (PTFE) thread sealant) to maintain solutions on surfaces during the reaction steps. All other surface modifications were performed directly in well plates. All incubation steps were performed in the dark with 90 rpm agitation on a rotary shaker (Ecotron, Infors HT) at room temperature. After each reaction step, solutions containing reactants were removed, and surfaces were rinsed twice with 0.2 μm-filtered PBS. Collagen-coated surfaces were used as a native protein control and were prepared by adding 0.15 mL/cm^2^ of 50 μg/mL type 1 rat-tail collagen (Thermo Fisher Scientific™) in 0.02 N acetic acid (Thermo Fisher Scientific™). All surfaces were sterilized by 95% ethanol.

### 2.2 X-Ray Photoelectron Spectroscopy (XPS)

The chemical composition of the aminated surfaces, before and after activation with S-SMPB, was investigated by XPS using a PHI 5600-ci spectrometer (Physical Electronics, Eden Prairie, MN). The main XPS chamber was maintained at a base pressure of < 8×10^−9^ Torr. A standard aluminum X-ray (Al Kα = 1486.6 eV) source was used at 300 W to record survey spectra with charge neutralization. The detection angle was set at 45° with respect to the normal of the surface and the analyzed area was 0.5 mm^2^.

### 2.3 Amino Detection via the Orange II Assay

The surface concentration of primary amines was quantified using the Orange II assay^25, 26^. The Orange II dye has a negatively charged sulfonated group that can preferentially bind to positively charged protonated primary amines in an acidic solution. The dye can then be released by adjusting the pH and then colorimetrically quantified. Surfaces cut into 1 cm^2^ surface-modified polystyrene pieces were transferred into 10 mL polystyrene tubes (#T406-2, Simport scientific, Beloeil, Ca). The samples were then submerged in Orange II sodium salt solution (14 mg/mL Orange II, 75370, Sigma Aldrich in RO water adjusted to pH 3 with 37% HCl; 1.5 mL added per tube). After incubating for 30 min at 40°C, samples were rinsed with the acidic solution to remove all unbound dye and air-dried before being immersed in 1 mL of alkaline solution (RO water adjusted pH to 12 with 5N NaOH solution) to desorb the dye. The pH of the desorbed dye solution was then readjusted to a pH of 3 by adding 1% v/v of 37% HCl to each tube. All solutions were transferred to a cuvette and absorbance measurements were taken at 484 nm on a Genova spectrophotometer (Jenway, Staffordshire, UK). Primary amines were quantified by comparing the absorbance obtained to a standard curve generated by adding Orange II dye in acidic solution at known concentrations ranging between 0.3 μg/mL and 140 μg/mL.

### 2.4 Contact Angle Measurements

The contact angles between deionized RO water and functionalized surfaces were measured by the sessile drop method using an OCA 150 system (DataPhysics Instruments GmbH, Filderstadt). Water drops of 5 μL were deposited at a rate of 0.5 μL/s onto Purecoat ™ substrates, with or without S-SMPB treatment. Images of the drops in contact with surfaces captured at the end of drop spreading were recorded. The average between the left and the right static contact angle values were determined for each image using the SCA-20 software (DataPhysics Instruments).

### 2.5 Enzyme-Linked Immunosorbent Assay (ELISA) for Protein G

A direct ELISA was developed to detect and quantify protein G surface concentrations. Purecoat™ 96 well plates (BD Purecoat™ Amine # 356717, BD Biosciences) were functionalized as described above (S-SMPB(+)) or by omitting the S-SMPB activation step (S-SMPB(−)). To block further protein adsorption, 200 μL/well of 1% BSA solution in PBS was introduced and left to incubate for 90 min at 37°C on a rotary shaker at 90 RPM. Wells were rinsed twice with washing buffer consisting of 0.05% Tween-20 (#P1379, Sigma-Aldrich) solution in PBS. To detect protein G, a chicken immunoglobulin Y (IgY) anti-protein G was used. This antibody was selected due to the absence of affinity between protein G and the Fc fragment of IgY antibodies: only the antigen binding fragment of the IgY anti-protein G can interact with protein G, which should facilitate quantification of surface ligands. A volume of 100 μL of horseradish peroxidase (HRP)-conjugated IgY anti-protein G secondary antibody (HRP anti-protein G, OAIA00498, Aviva systems biology) solution (0.02 μg/mL of HRP anti-protein G diluted in rinsing solution with 1% BSA) was incubated in each well for 2 h at room temperature. Wells were immediately rinsed once with 1% SDS-TRIS solution at pH 11 and twice with washing buffer. To detect HRP, 100 μL of Slow TMB-ELISA substrate solution (#34024, Thermo Fisher) was added per well. After 25 min of incubation without agitation at room temperature, 100 μL/well of 1M sulfuric acid solution was added to stop the reaction, and absorbance measurements were immediately taken at 450 nm on a Benchmark™ plate reader (Bio-Rad, Berkeley, USA).

### 2.6 Immobilized Antibody Detection and Quantification

Purecoat™ amine surfaces were modified as described above, except that only certain regions of test surfaces were treated with protein G by adding spots of 0.5 μL Cys-Protein G solution at concentrations ranging between 0.055 μM and 55 μM. To assess the effect of adsorption on surface amounts of protein G, the spots were deposited on surfaces with (S-SMPB (+)) or without (S-SMPB (−)) S-SMPB treatment. After 1 h incubation, surfaces were rinsed with PBS and covered (spot and surrounding region) with primary antibody solution for 1 h as described above. After two washes in PBS, surfaces were covered with Alexa Fluor 488-conjugated goat anti-mouse secondary antibody solution at 20 μg/mL. After 1 h of incubation at room temperature, surfaces were rinsed twice with 1% SDS-TRIS solution, twice with PBS and twice with RO water before air drying. Spots were then imaged using a laser scanning confocal microscope (Zeiss LSM 5 Exciter, Germany) at 10X with an argon laser (488 nm). A total of 10 images per spot were taken to obtain the mean fluorescence intensity of one spot along with the associated standard deviation value. A total of 3 spots per replicate were studied to obtain the mean fluorescence intensity of each condition.

### 2.7 ECFC Capture Under Flow

Peripheral blood mononuclear cells (PBMCs) were isolated from fresh adult human peripheral blood and ECFCs were expanded as previously described^27^. Fresh blood samples were collected from adult donors under informed consent following a protocol (Study No. A06-M33-15A) approved by the Ethics Institutional Review Board at McGill University. To study cell capture by antibody-modified surfaces under flow, functionalized surfaces were assembled into a custom parallel-plate flow chamber system with 4 independent chambers and flow paths, as previously described^28^. The flow chamber was sterilized and then assembled inside an incubator with humidified air maintained at 5% CO_2_ and 37 °C. Each chamber was connected to a reservoir that was pre-filled with 15 mL of warm serum-free EGM-2 (endothelial cell growth medium-2 without serum added from the kit, Lonza). ECFCs were harvested and resuspended in serum-free EGM-2 and added to the reservoirs to reach an overall cell density of 125,000 cells/mL. A peristaltic pump (Masterflex RK-7543-02 with Masterflex L/S two channels Easy Load II pump head using L/S 13 BPT tubing) was used to circulate the cell suspension in the system at a flow rate of 0.18 mL/s to obtain 1.5 dyn/cm^2^ wall shear stress. After 1 h of circulation, cells were fixed using a 4% paraformaldehyde solution (VWR) for 10 min, rinsed once in PBS and stored in PBS for immunocytochemistry.

### 2.8 Immunohistochemistry and Microscopy

Fixed cells were permeabilized for 15 min with 0.1% Triton X (VWR) in PBS. Nuclei were stained with 1 μg/mL DAPI (Sigma) diluted in RO water for 10 min. Slides were then rinsed with RO water and stored in PBS before being imaged on an inverted fluorescent microscope (Olympus IX81). Images were acquired at 10X in phase contrast and fluorescence. At least 40 phase contrast images were acquired on each slide per cell capture experiment. Captured cells were enumerated using the “analyze particles command” of ImageJ from the 40 acquired DAPI images.

### 2.10 Statistical Methods

Statistical analysis was performed with JMP Pro 13 software (SAS Institute, Cary, NC). Unless otherwise stated, data represent the average ± the standard deviation of 3 independent experiments. For water contact angle measurements, the reported values represent the average ± the standard error of the mean of 10 images per surface from 3 independent replicas. The criteria for statistically significant differences was selected to be p < 0.05. Student’s t-test was used for comparisons between two sample groups and comparisons between multiple groups were performed using two-way analysis of variance (ANOVA) followed by Tukey-Kramer HSD post hoc test. For ECFC capture experiments, each of the 4 replicas was conducted with ECFCs derived from a different donor.

## 2. Results

To develop a suitable antibody screening platform for cell capture, the proposed surface modification steps were first characterized, followed by testing the effect of different immobilized antibodies on ECFC capture under laminar flow.

### 3.1 Characterization of the Purecoat™ Substrate and S-SMPB Activation of the Surface

Cys-Protein G was conjugated onto commercially-available aminated polystyrene surfaces via the amine-to-sulfhydryl linking arm S-SMPB (Figure 1). The presence of S-SMPB on the surface prior to Cys-Protein G conjugation was assessed by studying the atomic composition of the surface by XPS. As expected, the nitrogen content decreased after S-SMPB treatment (Figure 2A). The carbon content increased, and oxygen content decreased after the S-SMPB reaction due to the elemental composition of the linking arm. The surface density of amino groups, based on the concentration of surface-bound Orange II, decreased after the S-SMPB activation step and was significantly higher on aminated Purecoat™ substrates compared to the control polystyrene surfaces.

**Figure 2:**
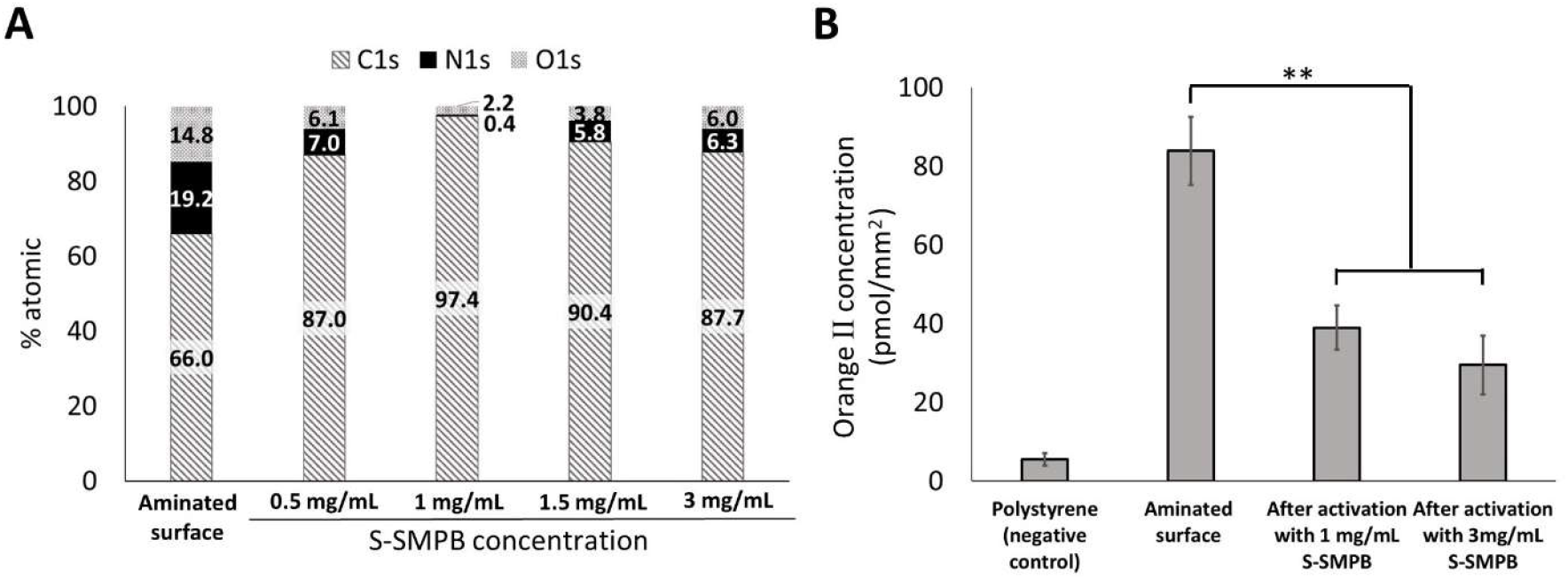
Characterization of the amine content on aminated polystyrene substrates before and after S-SMPB activation. (A) Summary of X-ray photoelectron spectroscopy measurements of the aminated substrate before and after activation with different concentrations of S-SMPB. Full set of data ± standard deviation are available in Table S.1. (B) Amino quantification of substrates using Orange II dye before and after activation with S-SMPB. Untreated polystyrene was used as a negative control. **P < 0.01 with N=3

Significant changes in surface amino content and presence of maleimide groups after S-SMPB activation are expected to significantly alter the surface free energy of the substrates which can be quantified through surface wettability. The static water contact angle with aminated and S-SMPB activated surfaces provides information about changes in the surface wettability due to surface modification. As shown in Figure 3, the reaction of S-SMPB with aminated surfaces for 2 h at 1 mg/mL or 3 mg/mL increased the contact angle. The decreased surface hydrophilicity observed after S-SMPB treatment is consistent with the decreased surface density of hydrophilic amino groups previously observed by XPS and addition of hydrophobic moieties present in the S-SMPB structure. Results from the XPS, Orange II assay and contact angle measurements are consistent with robust surface activation via S-SMPB applied at 1 mg/mL to 3 mg/L for 2 h.

**Figure 3:**
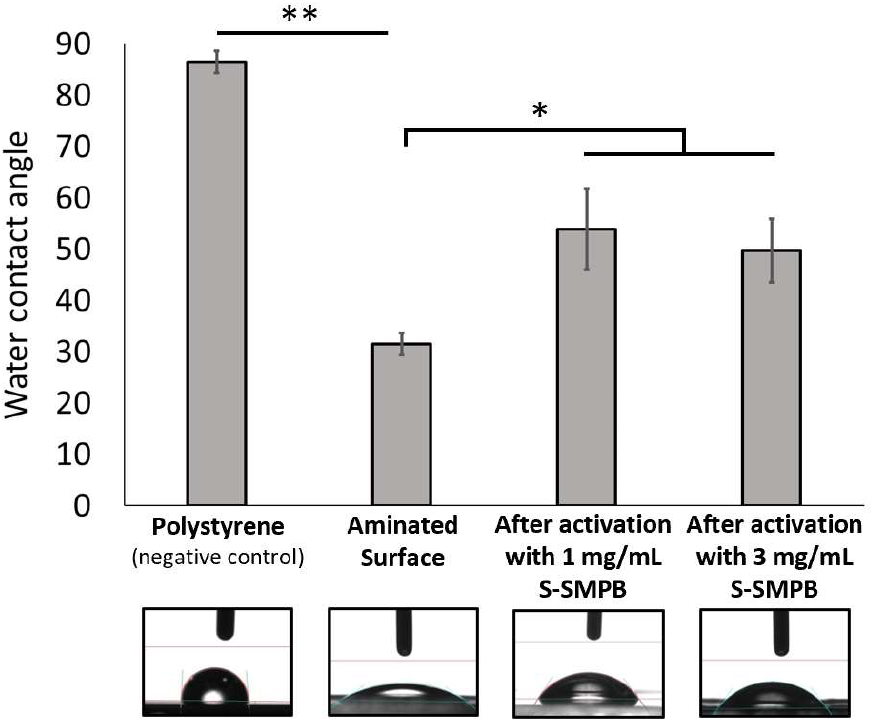
Water contact angle on aminated polystyrene substrates before and after activation with S-SMPB. S-SMPB was applied at 1 mg/mL and 3 mg/mL prior to rinsing, drying and goniometry. *P < 0.05; **P < 0.01 with N=3

### 3.2 Analysis of Protein G Grafting Efficiency

Next, Cys-protein G grafting was evaluated with a direct ELISA developed to detect and confirm protein G presence on surfaces (Figure 4). As shown in Figure 4, a positive correlation between protein G concentration and absorbance signal was observed under covalent conjugation conditions. This correlation was not observed for adsorbed Cys-Protein G in the absence of the linking arm. This suggests that the covalent conjugation method via S-SMPB activation improved control over the amount of Cys-Protein G present on surface compared to adsorption. Together, the results shown in Figure 2, Figure 3, and Figure 4 highlight the effectiveness of this covalent conjugation strategy in grafting Cys-Protein G in controlled amounts on amine-functionalized surfaces.

**Figure 4:**
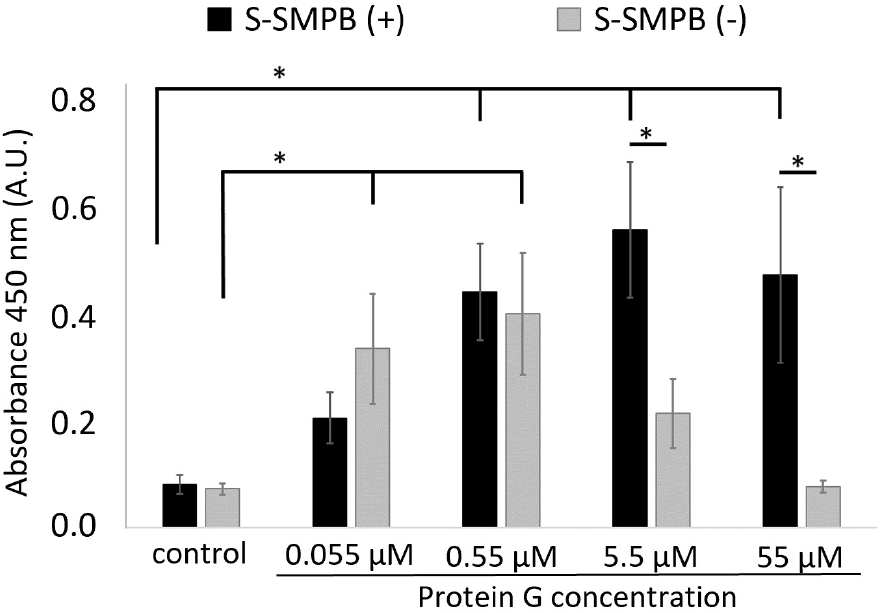
Direct ELISA detection of surface-immobilized protein G after covalent conjugation or adsorption (S-SMPB omitted from the reaction steps) while varying the concentration of protein G in solution. Control: no protein G added. *P < 0.05 with N = 3.

### 3.3 Antibodies Interact Specifically with Protein G Treated Surfaces

Having achieved covalent conjugation of protein G, the next step was to immobilize IgG antibodies onto the functionalized surfaces. As shown in Figure 5B, anti-CD31 antibodies were successfully immobilized on surfaces functionalized with protein G based on fluorescent secondary antibody detection. The fluorescence intensity in the region where protein G was deposited was significantly higher when applying 5.5 μM of protein G as compared to 0.55 μM or to the surrounding region without protein G. This was not observed on surfaces with adsorbed protein G (S-SMPB(−)). These experiments were repeated with four different IgG antibodies targeting endothelial (anti-CD31, anti-CD105, anti-CD144) or macrophage/monocyte (anti-CD14) surface markers which were successfully immobilized on surfaces using the same strategy (Figure 5C).

**Figure 5:**
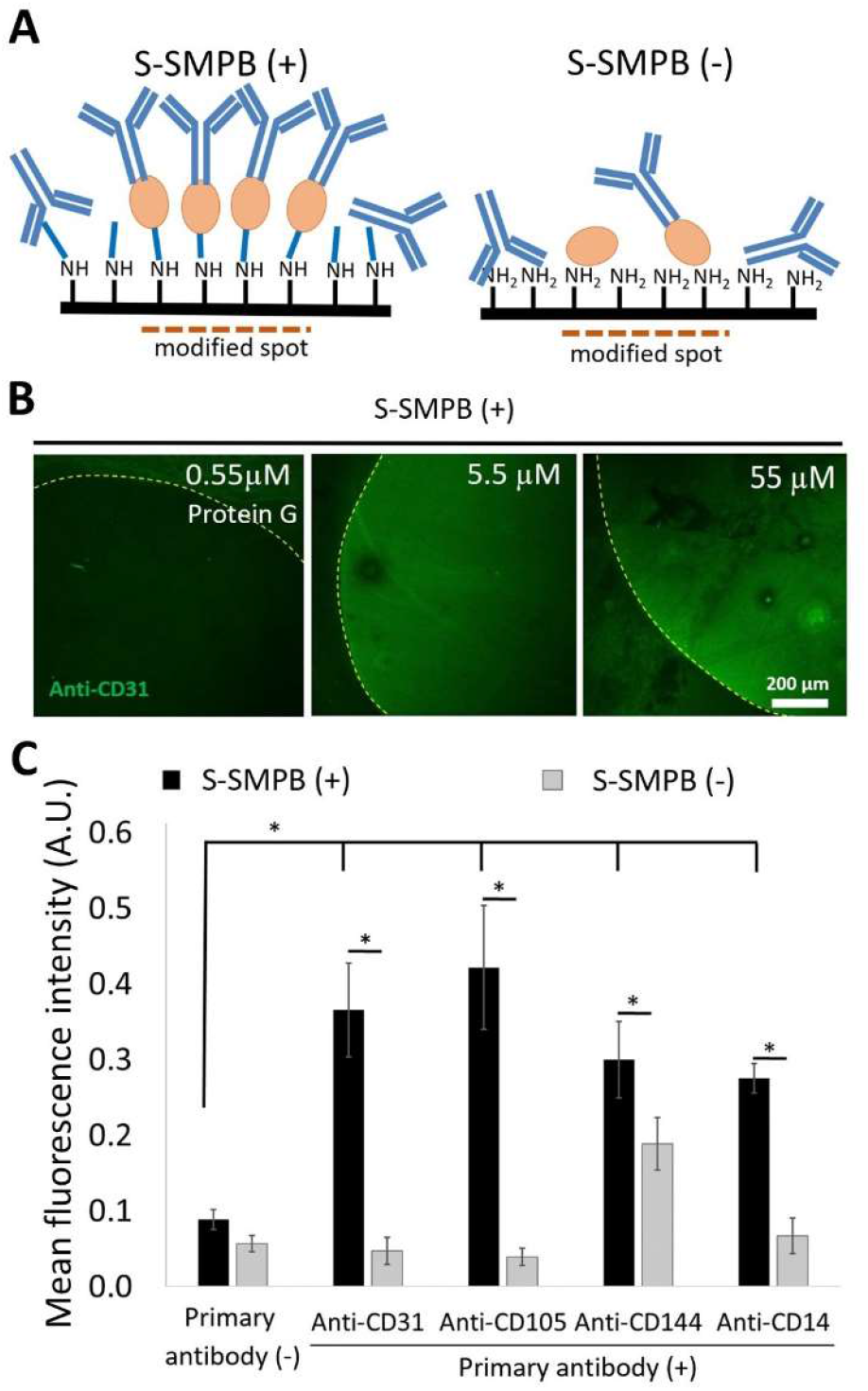
Fluorescence-based detection of immobilized IgG antibodies on conjugated protein G spots. (A) Schematic representation of antibody immobilization on protein G spots with (S-SMPB(+)) or without (S-SMPB(−)) covalent grafting of Protein G. (B) Fluorescence images of the protein G modified spots (at different protein G concentrations) that were then treated with primary anti-CD31 antibody and a fluorescently tagged secondary antibody. (C) Successful immobilization of different primary antibodies on conjugated protein G (S-SMPB (+)) based on the detection of fluorophore-labeled antibodies added after protein G and primary antibody immobilization. Surfaces without S-SMPB and/or without primary antibodies were used as negative controls. *P < 0.05 with N = 3.

### 2.4 Antibody-Functionalized Surfaces can Capture ECFCs

ECFCs were injected into a flow loop with a parallel plate chamber and circulated for 1 h at 1.5 dyn/cm^2^ wall-shear stress to determine whether antibody-modified surfaces can mediate cell capture. This wall shear stress is at the lower end of the physiological range and was selected to allow quantification of cell capture *in vitro*^29, 30^. Surfaces with (1) adsorbed anti-CD144 (S-SMPB (−)), (2) collagen-coated surfaces, (3) immobilized anti-CD14 on conjugated protein G (S-SMPB (+)) and (4) immobilized anti-CD144 on conjugated protein G (S-SMPB (+)) were tested in the flow system. Out of the four conditions, only the immobilized anti-CD144 on the conjugated protein G had a significant effect in enhancing ECFC capture under flow (Figure 6). Surfaces with adsorbed anti-CD144 showed no significant difference in the number of captured ECFCs compared with surfaces modified with anti-CD14, a surface antigen that is not expressed by ECFCs^27^. Surfaces coated with rat tail collagen, a commonly used ECFC substrate, also had a significantly lower number of captured cells compared to the immobilized anti-CD144 on the conjugated protein G. As a control, PBMCs, rich in CD14+ cells (>45%) but with low or undetectable CD144+ cell populations (<0.1%), were separately circulated in the same conditions over the same surfaces. In this arrangement, a significantly higher number of captured cells were observed on surfaces with immobilized anti-CD14 compared to surfaces with anti-CD144, confirming the specificity of the cell capturing strategy (Figure S.1).

**Figure 6:**
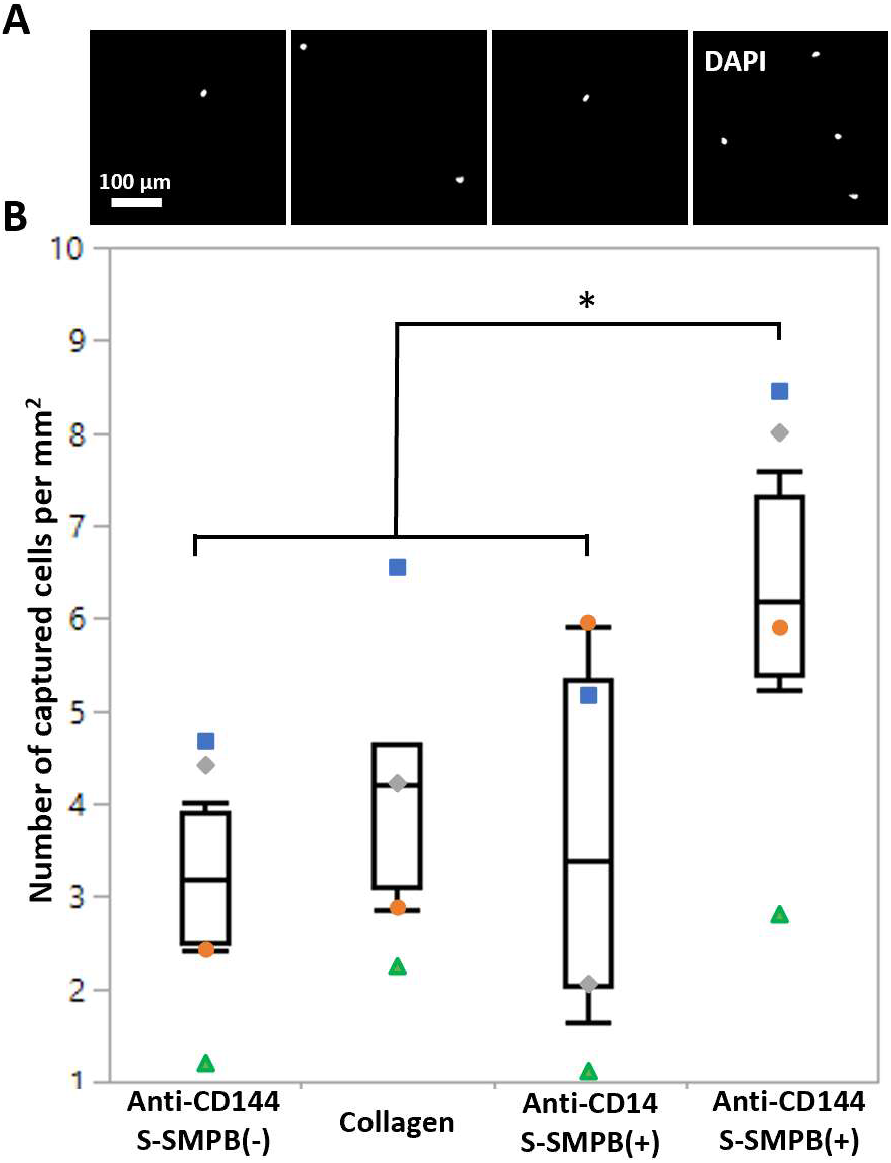
ECFC capture from laminar flow conditions over either anti-C144 (present on ECFCs) on adsorbed or conjugated protein G, anti-CD14 (not present on ECFCs) on conjugated protein G (negative control), or collagen. A) Fluorescence images of ECFC nuclei on modified surfaces after 1 h of exposure to cell suspension under flow conditions. B) Quantification of number of cells per mm2 on the modified surfaces at the end of the 1 h of flow. Each symbol represents data collected using ECFCs from a separate donor. *P < 0.05 with N = 4.

## 3. Discussion

To our knowledge, this study in the first demonstration of the utility of the covalent conjugation of Protein G, a molecule that is heavily relied upon for the production of biotherapeutics, in effectively immobilizing capture antibodies to create surfaces with enhanced ECFC cell capture potential. Our proposed oriented antibody surface modification strategy consists of immobilizing antibodies via the Fc region to Cys-Protein G that is conjugated to an aminated surface via a S-SMPB heterobifunctional linking arm. This strategy was selected to maximize the immunoaffinity of the antibodies compared to adsorption or direct covalent conjugation methods which can result in random orientation and reduced availability of antigen-binding sites^31–33^. The mild conditions (physiological pH, room temperature, aqueous conditions) of this surface modification strategy assures compatibility with a wide variety of cell culture substrates and biomaterials. The covalent conjugation of the protein G via its cysteine tag enhances the ability to control the orientation of the protein G and of the immobilized antibody^24^.

Different IgG antibodies were successfully immobilized on conjugated protein G, achieving better control over protein G surface density compared with adsorption (omission of the linking arm). The surfaces with grafted protein G and immobilized anti-CD144 successfully captured circulating ECFCs at 1.5 dyn/cm^2^ wall shear stress, contrary to surfaces where the linking arm was omitted from the surface treatment. The grafted protein G anti-CD144 surfaces also captured significantly higher circulating ECFC numbers compared with surfaces with antibodies which do not target ECFCs (anti-CD14). Compared to a native extracellular matrix protein such as collagen, the oriented surface-immobilized anti-CD144 antibodies resulted in significantly higher levels of ECFC capturing which demonstrates the significance of targeting specific cell-surface antigens. These promising findings highlight the value of our proposed surface modification strategy for the design of EPC capture vascular biomaterials.

S-SMPB is a versatile linking arm which has been applied to vascular biomaterials such as aminated polytetrafluoroethylene (PTFE)^34^, poly (L-lactide (PLLA), poly (ɛ-caprolactone) (PCL)^35^ and other aminated model surfaces^36^. The maleimide functional group of the S-SMPB reacts with sulfhydryl groups which is only found in the cysteine tag of the Cys-Protein G molecule, thus creating a selective oriented conjugation strategy. Our results show that the bio-affinity-based conjugation led to a better control over protein G surface density compared to protein G adsorption. With covalent Protein G grafting, surface concentration of protein G followed an expected saturation profile (Figure 4). Conversely, using protein G adsorption, the maximum achievable protein G surface concentration was lower and was followed by a decreased in surface densities at higher concentrations. A possible explanation for the drop in protein G concentration in adsorption condition is the ability of free cysteines on Cys-protein G to interact in solution to form disulfide bonds which can produce dimers reducing effective interaction with the detection antibodies of the ELISA. Therefore, the conjugation scheme with the S-SMPB linking arm allowed better control over protein G surface density.

We have recently shown that antibodies immobilized through the Fc region by surface conjugated RRGW can selectively capture ECFCs from a mixture of cells under dynamic flow conditions^20^. A major obstacle hampering the use of protein G on implanted biomaterials such as stents is its unknown immunogenic profile and the possibility that it can provoke undesirable host immune responses^37^. The RRGW peptide, on the other hand, can pose a lower risk of triggering an immune response due to its small chemically defined structure and the absence of endotoxins due to chemical synthesis. Larger protein structures such as protein G can also be more susceptible to enzymatic and thermal denaturization compared to smaller peptides such as RRGW. A potential advantage of the protein G strategy over the previously proposed RRGW peptide are the existing protein G supply chains allowing its use in cGMP bioprocessing plants, which may facilitate its large-scale use in other biomedical and clinical applications^38^. Furthermore, the recombinant Cys-Protein G has 25 times the molecular weight of the RRGW peptide which can lead to 2 to 3 nm additional spacing between the antigen-binding site and the modified surface, reducing steric hindrance^23^. There are three Fc fragment binding sites available on each protein G molecule compared to the RRGW peptide’s single antibody binding capacity, potentially increasing the density of antibodies which can be immobilized on protein G modified surfaces. Further development of antibody immobilization strategies via protein G and RRGW for *in vivo* use will require side-by-side comparison of the hemocompatibility, immunogenicity, and stability of both molecules. All in all, the presented analysis of the proposed antibody immobilization strategy provides a promising prospect for producing clinically successful EPC-capturing biomaterials.

## 4. Conclusion

This study presents a 3-step surface functionalization strategy to immobilize antibodies on aminated surfaces via Fc region interactions with Protein G. This technology can be applied to engineer endothelial progenitor cell capture stents and other cell separation devices. Model aminated polystyrene surfaces were first reacted with an amine to sulfhydryl linking arm. The linking arm was then used to conjugate protein G to the surface through a cysteine tag maximizing its antibody immobilization capacity. Different IgG antibodies were successfully immobilized on the surface and detected using a simple fluorescence-based approach. Finally, surfaces modified with anti-CD144 via our protein G-based approach displayed superior ability in capturing human derived ECFCs from flow compared to surfaces modified with passive adsorption. Our work highlights the potential of grafted protein G-based surface functionalization strategies in enhancing the potential of ECFC capture on the surface of vascular implants. Orienting the antibodies on EPC capture stents may accelerate the endothelialization process essential in vascular regeneration and homeostasis.

## Author Contributions

MB: conceptualization of initial hypothesis, design of experimental methodology, experimental investigation focused on surface characterization, data analysis and curation, writing, reviewing, and editing the manuscript.

MAE: design of experimental methodology, experimental investigation focused on cellular responses, data analysis and curation, writing, reviewing, and editing the manuscript.

OSB: experimental investigations focused on cell capture studies, reviewing, and editing the manuscript.

GL: funding acquisition, design of the experimental methodology, data analysis, reviewing and editing the manuscript.

CAH: funding acquisition, supervision, conceptualization of initial hypothesis, design of experimental methodology, data analysis and curation, writing, reviewing, and editing the manuscript, and project administration.

All authors contributed to the article and approved the submitted version.

## Acknowledgements

The authors thank Ariane Beland, Stéphanie Vanslambrouck, Pascale Chevallier, Lisa Danielczak, Ranjan roy, Frank Caporuscio, Natalie Fekete and Gad Sabbatier for technical support and Raymond Tran for his help in reviewing the manuscript.This study was financially supported by the Canadian Institute for Health Research (CIHR, MOP 142285) and the Canadian Foundation for Innovation (CFI, project 35507). This research was undertaken, in part, thanks to funding from the Canada Research Chairs Program. This work was supported via travel awards and networking opportunities offered by ThéCell (The Quebec Network for Cell, Tissue and Gene Therapy), the Quebec Center for Advanced Materials (QCAM), PROTEO (The Quebec Network for Research on Protein Function), CMDO (the Cardiometabolic Health, Diabetes, and Obesity Research Network), and the MRM (McGill Regenerative Medicine) network.

## Supplementary Information

**Table S.1:** X-ray photoelectron spectroscopy measurements of the aminated substrate before and after activation with different concentrations of S-SMPB reported as the mean of 3 replicas ± the standard deviation.

| S-SMPB conc (mg/mL) | C1s | N1s | O1s |
| --- | --- | --- | --- |
| Aminated surface | 66.0 ± 1.0 % | 19.2 ± 0.9 % | 14.8 ± 1.0 % |
| 0.5 | 87.0 ± 3.2 % | 7.0 ± 2.0 % | 6.1 ± 1.2 % |
| 1 | 97.4 ± 1.1 % | 0.4 ± 0.4 % | 2.2 ± 0.9 % |
| 1.5 | 90.4 ± 1.7 % | 5.8 ± 0.7 % | 3.8 ± 1.0 % |
| 3 | 87.7 ± 0.6 % | 6.3 ± 0.2 % | 6.0 ± 0.6 % |

**Figure S.1:**
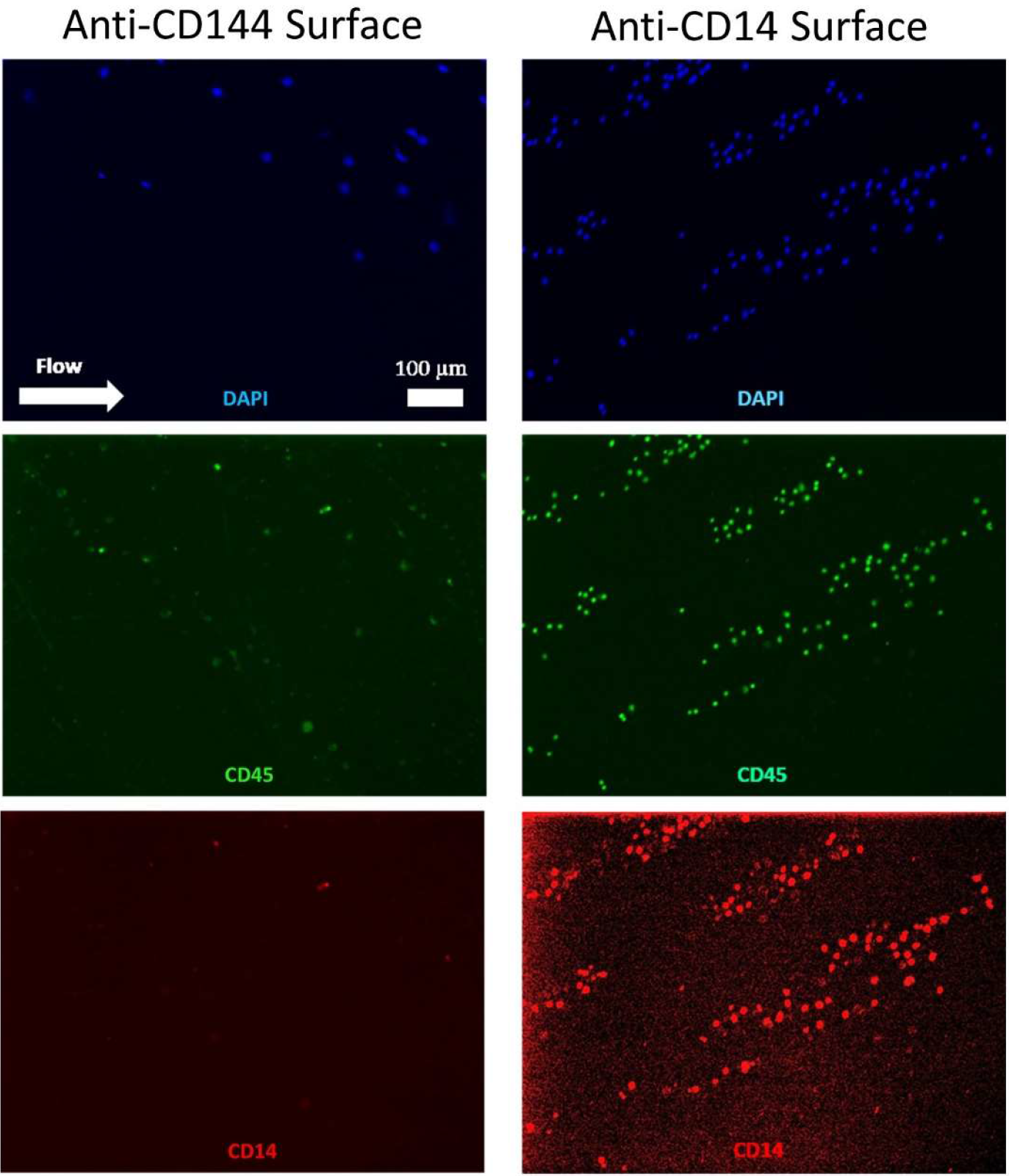
Representative images of PBMCs on surfaces with immobilized anti-CD144 or anti-CD14 antibodies on conjugated protein G. Nuclei (blue) were stained with DAPI, CD45 hematopoietic marker was detected with a rabbit anti-CD45 primary antibody and an AF488 goat anti-rabbit secondary antibody, and CD14 monocyte marker was detected with a mouse anti-CD14 primary antibody and an AF555 goat anti-mouse secondary antibody.

